# Monitoring Autonomic Tone During Spinal Cord Neuromodulation Using Wearable AURIS Sensor

**DOI:** 10.64898/2026.03.07.709943

**Authors:** Ryan S. Bohluli, Angelica F. Lopez, Pierce L. Perkins, Kenzi M. Griffith, Avaneesh Babu, Sung-Min Cho, Nitish V. Thakor

## Abstract

The transition of bioelectronic medicine to clinical use is currently limited by a lack of non-invasive sensors capable of measuring autonomic tone during active neuromodulation. Conventional monitoring modalities, such as mean arterial pressure (MAP) and Ag/AgCl chest electrodes, are often invasive, cumbersome, or susceptible to motion artifacts. Here, we present a novel framework employing an in-ear sensor (AURIS) to continuously monitor heart rate variability (HRV) during therapeutic neuromodulation. These sensors utilize a polydimethylsiloxane (PDMS) substrate to ensure biocompatibility and superior conformability. Experiments in a rodent model (*n = 3*) demonstrate that the AURIS platform achieves gold-standard fidelity, with mean heart rate differences of 6.03 BPM and mean RR interval deltas of 3.18 ms compared to chest electrodes. Sensor agreement was statistically validated using independent t-tests, showing no significant difference between modalities (all *p > 0.46*). While time-domain shifts trended toward significance, complexity metrics showed robust sequential responses with large effect sizes, including the SD1/SD2 ratio (*d = 1.474*) and the DFA α ratio (*d = 1.091*). These findings validate a sensor architecture that is durable, accessible, and provides the necessary technical foundation for closed-loop feedback and non-invasive clinical trials.

## I. INTRODUCTION

The integration of high-precision neuromodulation with wearable physiological sensors is a cornerstone of the emerging field of bioelectronic medicine. For patients with autonomic dysregulation, real-time monitoring of physiological parameters during non-invasive stimulation is necessary [1]. Traditional autonomic monitoring relies on mean arterial pressure (MAP) and heart rate variability (HRV) chest-based electrocardiogram (ECG) electrodes. However, MAP is often a lagging indicator of neural state and requires invasive arterial catheterization to achieve high temporal resolution [2] [3] [4]. HRV is derived from conventional ECG recordings by detecting the R-wave to calculate the beat-to-beat interval. The current gold-standard sensor for ECG recordings is Ag/AgCl chest electrodes, which have significant limitations. These sensors require electrolytic gels that evaporate over time, utilize adhesives that can cause skin irritation, and are highly susceptible to motion artifacts during active stimulation [5]. There is a critical need for a sensor platform that delivers gold-standard fidelity while maintaining the durability and accessibility of a wearable device. Here, we present our novel sensing and modulation framework, AURIS, combining a flexible, in-ear PEDOT:PSS conductive polymer sensor with a method for recording high-fidelity ECG signals to measure autonomic tone. The AURIS sensor platform leverages the stable, low-impedance environment of the auditory canal to achieve a signal-to-noise ratio comparable to that of conventional chest electrodes while minimizing motion-induced noise [6]. This setup is also resistant to other auditory confounds through the blocking of the ear canals, such as the “startle response” to ultrasound, which often plagues traditional ECG setups [7]. Focused ultrasound (FUS) is a precise tool for cardiac autonomic modulation. However, its clinical translation is hindered by a lack of robust, non-invasive sensors capable of real-time quantification of autonomic function [8] [9]. Building on previous demonstrations that thoracic FUS reduces MAP, this study aims to validate these effects through the continuous monitoring of HRV [4]. This work validates AURIS sensors as a viable alternative to the clinical gold standard electrodes. Using previously established surgical parameters [10] [11], we can show that the AURIS sensor effectively captures the same clinically relevant parameters as invasive chest leads with a minimal delta.

## II. METHODS AND MATERIALS

### A. Sensor Manufacturing

To capture high-fidelity ECG data to derive HRV, two custom PEDOT:PSS-based sensors were fabricated following a methodology from previous literature [6]. The sensors consist of a polydimethylsiloxane (PDMS) and PEDOT:PSS blend. This solution is dispensed into an SDS-treated, pipette-tip cast to form the final AURIS electrode. This electrode was made compatible with a blood pressure monitoring system (**Fig. 1**).

**Fig. 1.**
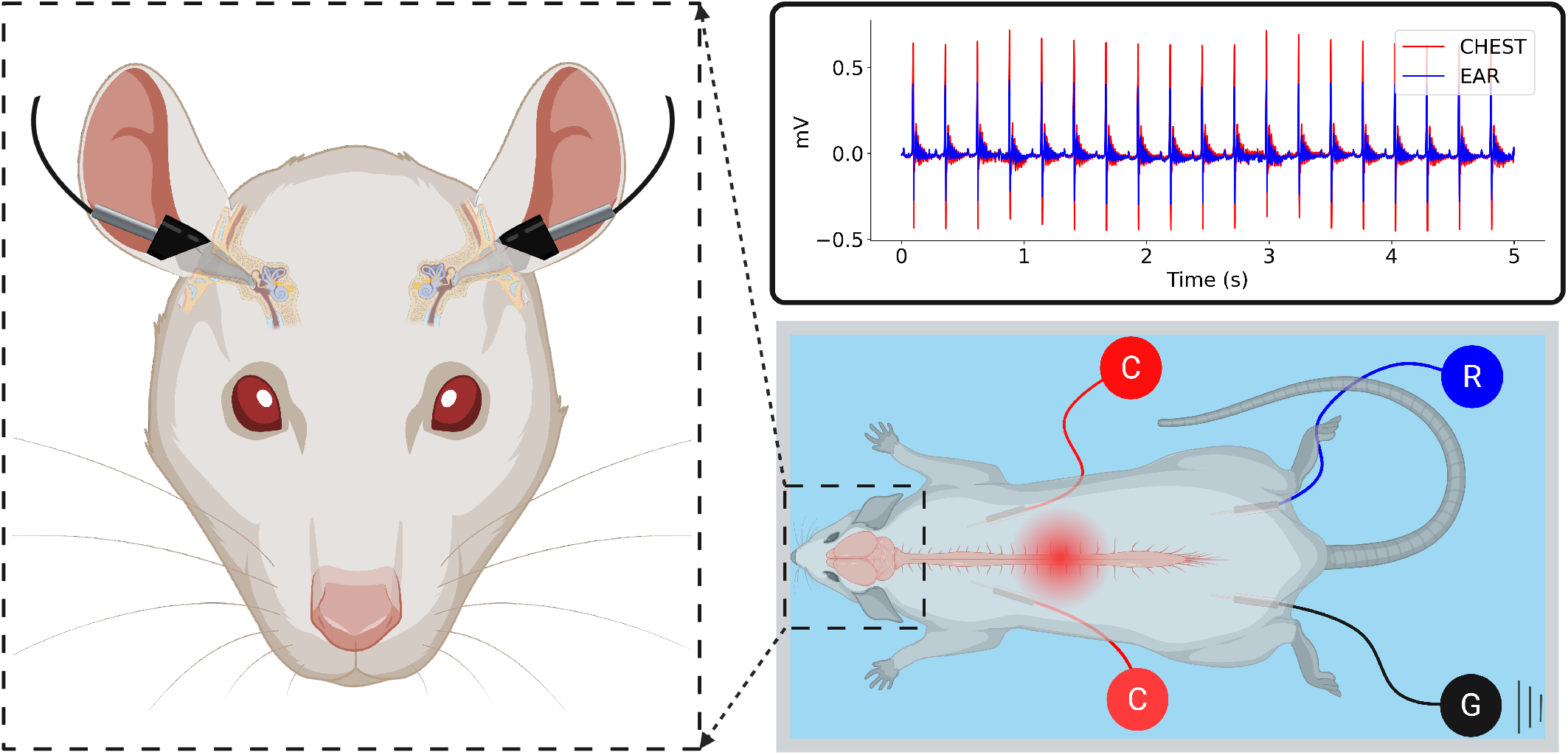
Experimental setup for autonomic monitoring. Schematic of bilateral AURIS in-ear electrodes recording ECG. Bilateral chest electrodes (C) corresponding to lead I (Enthoven) recording gold standard ECG for comparison. Bilateral thigh electrodes recording ground (G) and reference (R) signals. The graph displays the in-ear and chest ECG signals simultaneously.

Before placement within the auditory canals, the sensors were filled with conductive gel (Bio-Medical Instruments) to minimize resistance and noise [4] [12]. These were supplemented by two surface electrodes implanted into the chest. A ground electrode and a reference electrode were placed in the posterior thigh muscle (**Fig. 1**).

ECG signals were acquired and digitized using a Tucker-Davis Technologies (TDT, Alachua, Florida) data-acquisition system (RZ5D Real-Time Processor). The data was recorded, filtered, and monitored in real-time using TDT’s Synapse software. This setup ensured high-fidelity recordings (1 kHz sampling rate) necessary for accurate R-peak detection in rodents [13].

### B. Surgical Setup

This study involved adult female Sprague-Dawley rats (*n = 3*) for all procedures. Rats were anesthetized with isoflurane, initially induced at 2.7-3.0% and reduced to 1.8-2.2% during surgery. Invasive blood pressure was monitored via a 30-G catheter inserted into the femoral artery and connected to a blood pressure monitoring system. Two methods were used to ensure proper euthanasia: the inhaled carbon dioxide method and confirmation through cervical dislocation. All experiments followed the guidelines of the National Institute of Health for laboratory animal care, and protocols were approved by the Johns Hopkins University Animal Care and Use Committee [4] [10].

The rats in the study had FUS stimulation delivered either invasively or noninvasively. Invasive surgery involved a standardized procedure for a lower laminectomy to expose the spinal cord [3]. Anesthesia was maintained via monitored isoflurane adjustments. Depth was confirmed using the paw pinch reflex and by maintaining a respiratory rate of approximately 60 breaths per minute. For invasive trials, a T12–T13 laminectomy was performed to expose the thoracic spinal cord for direct sonication. Non-invasive subjects received a midline skin incision over the T13 vertebra, leaving the underlying musculature and bone intact [10].

After establishing the surgical environment, FUS was administered to the thoracic spinal cord at T12-T13 in stepwise increases. The main parameters for coaxial modulation were voltage amplitude (mVpp), stimulation frequency (kHz), stimulation duration (seconds), and duty cycle (%). To reduce variability among the rats, the ultrasound parameters were fixed at 500 kHz and 250 mVpp [10].

### C. Autonomic Tone Monitoring

A custom signal-processing pipeline in Python was developed to detect R-R intervals [6]. To ensure robust detection, R-R intervals were extracted independently from both the AURIS and the chest electrode channels (Enthoven lead I). These independent signals were compared and averaged to generate a single, aggregate R-R time series for the final analysis (**Fig. 3**). This multi-lead integration minimized the impact of local motion artifacts and ensured high-fidelity HRV data.

The Python pipeline analyzed the R-peaks and produced several outputs: a 5-second snippet of the normalized and denoised ECG data (**Fig. 1**), a tachogram illustrating the HRV trends throughout the experiment, Poincaré plots for both the stimulation and the washout period displaying the distribution of all HRV values, and a power spectral density graph for the stimulation and washout periods. The pipeline was designed to output a comprehensive summary of physiological parameters for comparative analysis.

To quantify the magnitude of physiological shifts induced by neuromodulation, Cohen’s *d* and a paired t-test were calculated for each HRV metric, comparing the stimulation and subsequent washout periods. Cohen’s *d* was the primary metric for evaluating statistical power, with values exceeding 0.8 considered large effects and values exceeding 0.5 considered medium effects, which are deemed sufficiently powerful for the purposes of this experiment. Sensor modality agreement was validated using independent t-tests to ensure cross-platform fidelity during these high-power transitions.

All data generated in this study will be made available on request. For data, please contact Ryan Bohluli. The code used in this manuscript for statistical analysis is available on GitHub at the following domain: https://github.com/afl008/AURIS.

## III. RESULTS AND DISCUSSION

### A. Sensor Agreement

The primary objective was to validate the AURIS sensor against the established gold standard of chest-based Ag/AgCl electrodes. **Table 1** summarizes baseline metrics across all subjects (*n = 3*). These include time-domain (e.g., Mean RR, RMSSD), frequency-domain (e.g., Total Power), and non-linear complexity measures (e.g., Sample Entropy). Across all subjects, the raw values recorded by the AURIS sensor closely mirrored those of the chest leads (**Table 1**).

**Table 1.** Physiological Parameters.

|  | Rat 1 |  | Rat 2 |  | Rat 3 |  |
| --- | --- | --- | --- | --- | --- | --- |
| Site | *Chest* | *Ear* | *Chest* | *Ear* | *Chest* | *Ear* |
| Mean RR | 149 | 146 | 221 | 221 | 145 | 146 |
| Mean HR | 451 | 430 | 277 | 277 | 414 | 411 |
| SDNN | 6.14 | 6.05 | 4.34 | 4.28 | 16.6 | 16.7 |
| RMSSD | 7.74 | 6.86 | 4.67 | 4.55 | 23.9 | 23.6 |
| PNN5 | 25.3 | 22.27 | 23.8 | 23.9 | 83.8 | 83.4 |
| LF | 7.36 | 6.66 | 5.96 | 5.90 | 52.8 | 52.9 |
| HF | 15.8 | 11.4 | 5.13 | 4.92 | 118 | 114 |
| Total Power | 23.2 | 18.1 | 11.1 | 10.8 | 171 | 167 |
| LF/HF | 0.474 | 0.581 | 1.76 | 2.06 | 0.448 | 0.460 |
| SD1 | 5.48 | 4.85 | 3.30 | 3.22 | 16.9 | 16.7 |
| SD2 | 6.72 | 6.95 | 5.12 | 5.08 | 16.4 | 16.7 |
| SD1/SD2 | 0.835 | 0.751 | 0.638 | 0.632 | 1.03 | 0.999 |
| Sample Entropy | 0.962 | 0.891 | 1.67 | 1.71 | 2.15 | 2.21 |
| DFA $\alpha 1$ | 0.814 | 0.875 | 0.719 | 0.722 | 0.613 | 0.641 |
| DFA $\alpha 2$ | 0.694 | 0.742 | 0.613 | 0.622 | 0.508 | 0.498 |
| DFA $\alpha$ Ratio | 1.23 | 1.21 | 1.75 | 1.74 | 1.22 | 1.34 |

The recorded parameters included the time difference between R-peaks, measured in milliseconds, and the mean heart rate, measured in beats per minute. These metrics were used to quantify changes in autonomic tone using established clinical measures [14].

The root mean square of successive differences (RMSSD) reflects the beat-to-beat variability in heart rate. Another metric, the percentage of adjacent normalized beat-to-beat (NN) intervals that differ by more than 5 milliseconds (PNN5), is also used to measure beat-to-beat variability, with the 5 ms threshold chosen to capture individual heartbeats in rats, whose typical heart rates range from 200 to 500 beats per minute. The standard deviation of NN intervals (SDNN), represents the deviation of those normal-to-normal beats.

Total power represents the sum of energy in the low-frequency (LF) and high-frequency (HF) bands, corresponding to the baroreflex and parasympathetic components of sympathovagal balance, respectively. The parameters SD1 and SD2 assess short-term vagal tone and long-term regulation, respectively. The sample entropy calculation represents autonomic complexity. Finally, detrended fluctuation analysis (DFA) α1 and α2 provide information about short-term and long-term autonomic scaling, respectively [14].

This agreement was statistically confirmed through independent t-tests, as shown in the “Modality Agreement” column of **Table 2**. No statistically significant differences were found between sensor sites for any metric (*p > 0*.*46* in all cases). Specifically, Mean RR (*p =* 0.834) and Mean HR (*p =* 0.771) demonstrated high cross-platform fidelity. These findings are visually supported by **Fig. 2** and **Fig. 3**, which show that both normalized HR distributions and the geometric patterns of Poincaré plots are virtually indistinguishable between the ear and chest sites.

**Table 2.** Sequential Analysis.

| | Modality Agreement (p-value)<sup>a</sup> | Sequential Effect (Mean $\Delta$)<sup>b</sup> | Cohen's $d$ (Power) | Sequential Significance (p-value) | |
| --- | --- | --- | --- | --- | --- |
|  |  |  |  | *Chest* | *Ear* |
| Mean RR | 0.834 | -3.18 | 0.821** | 0.029* | 0.064 |
| Mean HR | 0.771 | +6.03 | 0.746** | 0.043* | 0.093 |
| SDNN | 0.969 | -1.19 | 0.506** | 0.144 | 0.024* |
| SD1/SD2 Ratio | 0.992 | +1.07 | 1.474** | 0.007* | 0.122 |
| DFA $\alpha$ Ratio | 0.464 | +0.101 | 1.091** | 0.007* | 0.016* |

<sup>a</sup> Independent t-test comparing Chest vs. Ear modalities across all trials.
<sup>b</sup> Paired t-test comparing Stimulation vs. subsequent Washout periods (S-W).
\* Statistically significant ( $p < 0.05$ ) values.
\*\* Sufficiently powerful ( $d > 0.5$ ) values.

**Fig. 2.**
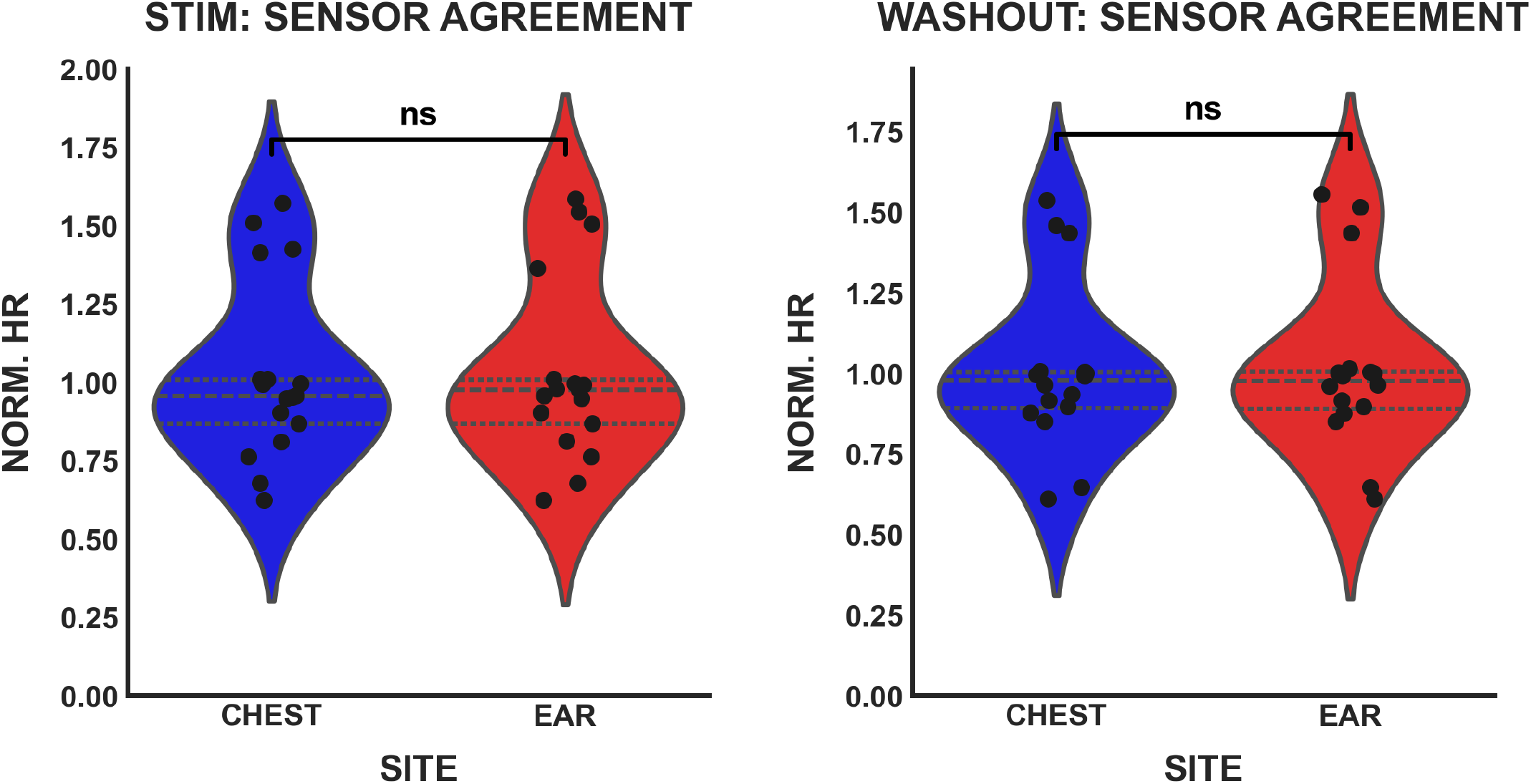
Violin plots demonstrating a comparison between normalized HR. HR was recorded from both sensors during stimulation (left) and washout (right) periods, with neither showing a statistically significant difference (*p > 0*.*05*).

**Fig. 3.**
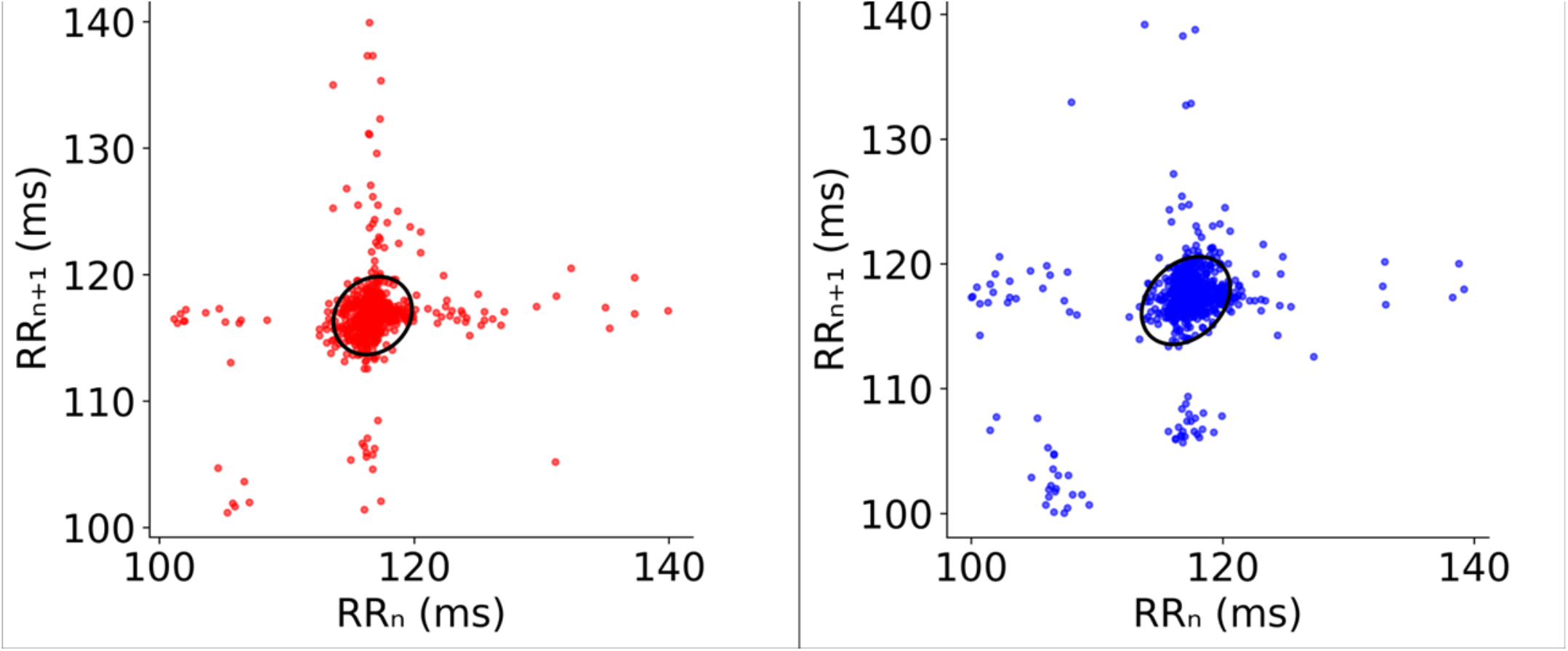
Comparison of Poincaré plots. Poincaré plots for a single FUS trial between the aggregated signal from chest electrodes (left) and the aggregated signal from in-ear electrodes (right).

### B. Sequential Autonomic Response to FUS

Sequential analysis between stimulation and subsequent washout periods revealed a detectable autonomic response to thoracic FUS. As detailed in **Table 2**, we observed a significant increase in the Mean RR interval (-3.18 ms, *p =* 0.029) and a corresponding decrease in Mean HR (+6.03 BPM, *p =* 0.043) within the chest recordings immediately following stimulation cessation.

While the AURIS sensor followed these physiological trends, the in-ear modality did not reach the *p < 0*.*05* significance threshold. For these time-domain metrics, the shifts trended toward significance at *p = 0*.*064* and *p = 0*.*093*, respectively. This discrepancy may be attributed to the small sample size (*n =* 3), as the magnitude of these transitions was nevertheless validated by high effect sizes. Complexity metrics demonstrated the most robust sequential shifts: the SD1/SD2 ratio showed the largest overall effect size (*d =* 1.474), while the DFA α ratio exhibited a significant shift during the washout transition (+0.101, *p =* 0.007) with a large Cohen’s *d* of 1.091.

While the sample size is limited, the high effect sizes observed across several metrics provide strong evidence of the platform’s sensitivity.

### C. Stability and Clinical Utility

Beyond statistical agreement, the AURIS system demonstrated practical stability over multiple stimulation trials. **Fig. 4** illustrates that the in-ear system provides accurate, continuous data that mirrors the chest signal’s power and trends throughout the experiment. Notably, the AURIS sensor’s design, leveraging a PDMS-based PEDOT:PSS substrate, addresses the critical limitations of traditional electrodes. Traditional Ag/AgCl leads suffer from motion artifacts and signal degradation due to gel evaporation. Conversely, in-ear sensors are decoupled from thoracic mechanical vibrations, which often compromise traditional ECG recordings [5].

**Fig. 4.**
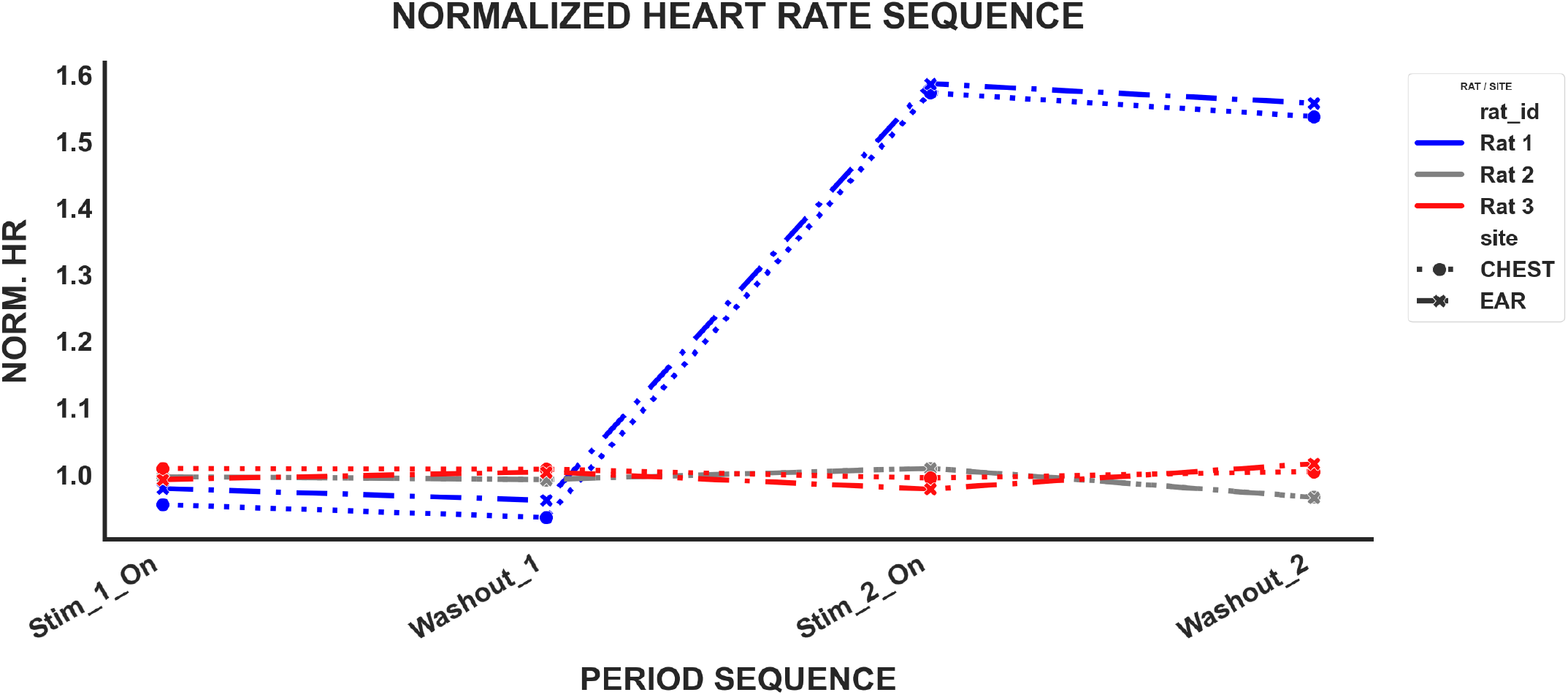
Performance of chest and ear electrode over time. Line graph illustrating the change in normalized heart rate (HR) over the course of multiple focused ultrasound (FUS) stimulation and washout trials for all three rats, showing the combined in-ear and chest signals for comparison.

By demonstrating that the in-ear platform captures high-complexity autonomic shifts with gold-standard fidelity, these results validate AURIS as a viable, less-invasive alternative for real-time cardiovascular monitoring during therapeutic neuromodulation. Limitations may include the possibility of sensors loosening and falling out of the ear canal, which is exacerbated with the use of conductive gel. For Rat 1, the aggregated signal was biased for portions of the recording due to one sensor falling out. We accounted for this by using only the signal from one AURIS sensor for that portion of the recording. This architecture addresses the clinical need for sensors that maintain high signal-to-noise ratios while remaining resistant to the motion artifacts and adhesive limitations common in traditional thoracic leads.

### D. Future Directions

The validation of the AURIS platform provides the necessary sensing infrastructure to transition from invasive animal studies toward non-invasive human clinical trials. To account for potential loosening of the sensors, the AURIS form factor can be molded to replicate the specific architecture of each patient’s ear canal. Future research should examine how AURIS can be integrated into closed-loop neuromodulation systems. In these systems, real-time HRV metrics could be used to dynamically adjust stimulation parameters, rather than relying on invasive metrics such as MAP. Ultimately, this framework enables a shift toward patient-centric bioelectronic medicine, facilitating continuous autonomic monitoring with minimal clinical burden during therapeutic interventions.

## IV. CONCLUSION

This study validates the AURIS sensor as a reliable, non-invasive alternative for monitoring autonomic tone during spinal cord neuromodulation. The platform achieves gold-standard fidelity, with a mean heart rate difference of 6.03 BPM and a mean RR interval delta of 3.18 ms compared to chest-based leads. Independent t-tests revealed no statistically significant difference between modalities across all recorded metrics (all *p > 0*.*46*). Although time-domain shifts recorded by the AURIS sensor trended toward significance (*p = 0*.*064*), it captured the magnitude of autonomic transitions through high-power complexity metrics. The SD1/SD2 ratio (*d = 1*.*474*) and DFA α ratio (*d = 1*.*091*) reliably detected shifts in sympathovagal balance across both invasive and non-invasive thoracic FUS interfaces. The AURIS sensor’s PDMS-based architecture addresses clinical barriers like gelation and motion artifacts while providing a reliable non-invasive correlate to hemodynamic changes in FUS studies. This validation enables larger sample sizes and more varied stimulation parameters in future studies. It promotes a move from invasive animal models to non-invasive cardiovascular management in human clinical trials.

## ACKNOWLEDGMENTS

R.S.B. and K.M.G. acknowledge support from the Johns Hopkins University Provost’s Undergraduate Research Award (PURA) Fellowship. A.F.L. and P.L.P. acknowledge support from the National Science Foundation Graduate Research Fellowship Program (NSF GRFP): award DGE-2139757. N.V.T. acknowledges funding from NIH R01 HL139158 and R01 HL071568. The authors thank Jeeun Kang and Ananya Tandri for providing the ultrasound equipment used in this experiment. They also thank A. Daniel Davidar and Nicholas Theodore for their consultation and guidance on rodent neurosurgery.

